# DNA methylation adjusts the specificity of memories depending on the learning context and promotes relearning

**DOI:** 10.1101/060152

**Authors:** Stephanie D Biergans, Charles Claudianos, Judith Reinhard, C Giovanni Galizia

## Abstract

The activity of the epigenetic writers DNA methyltransferases (Dnmts) after olfactory reward conditioning is important for both stimulus-specific long-term memory (LTM) formation and extinction. It, however, remains unknown which components of memory formation Dnmts regulate (e.g. associative vs. non-associative) and in what context (e.g. varying training conditions). Here we address these aspects in order to clarify the role of Dnmt-mediated DNA methylation in memory formation. We used a pharmacological Dnmt inhibitor and classical appetitive conditioning in the honeybee *Apis mellifera*, a well characterized model for classical conditioning. We quantified the effect of DNA methylation on naïve odour and sugar responses, and on responses following olfactory reward conditioning. We show that (1) Dnmts do not influence naïve odour or sugar responses, (2) Dnmts do not affect the learning of new stimuli, but (3) Dnmts influence odour-coding, i.e. 'correct' (stimulus-specific) LTM formation. Particularly, Dnmts reduce memory specificity when experience is low (one-trial training), and increase memory specificity when experience is high (multiple-trial training), generating an ecologically more useful response to learning. (4) In reversal learning conditions, Dnmts are involved in regulating both excitatory (re-acquisition) and inhibitory (forgetting) processes.

## 1 Introduction

The ability of honey bees to learn and form memories has been described and investigated in depth for many years (Menzel, 2012). When bees forage they search for good food sources and memorize their features such as color, shape and smell (Menzel, 2012). Bees show flower constancy during foraging (Chittka et al., 1999) and remember the features of a food source. On the other hand, it is also essential for bees to be able to re-evaluate their behaviour, if a source no longer provides good quality food (Greggers and Menzel, 1993).

Thus, extinction (i.e. forgetting) and re-acquisition are equally important. Furthermore, the environments bees encounter are variable, e.g. due to the slightly different smell of two flowers of the same species. Therefore, their ability to generalize stimuli belonging to the same category (e.g. type of flower) is as important as the ability to discriminate distinct stimuli (Cheng, 2000; Shepard, 1987). These different aspects and demands of foraging are reflected in bees cognitive capacities, and have been well documented in free-flying bees (Menzel, 2012).

Bee memory formation can be studied under controlled conditions with the proboscis extensions response (PER) (Bitterman et al., 1983). In this assay, bees learn to associate an odour with a sugar reward, similar to the olfactory learning taking place when a bee collects nectar from a flower during foraging (Eisenhardt, 2014). Depending on the conditions used during training, the dynamics of memory formation differ; for example multiple, but not one, odour-sugar pairings cause a prolonged increase of protein kinase A (PKA) (Hildebrandt and Muller, 1995; Muller, 2000). This suggests that different molecular pathways and dynamics may underlie memory formation depending on the training conditions.

Both few training trials and short time-intervals between training trials are associated with a reduced stimulus-specific memory, i.e. stronger generalization to novel stimuli (Lefer et al., 2012; Perisse et al., 2009). Generalization is the cognitive counterpart to perceptual discrimination (Cheng, 2000). It is dependent on stimulus similarity (e.g. different hues of blue, compared to yellow), but additionally requires a cognitive categorization of stimuli, which is experience dependent (Wright et al., 2008).

Relearning (e.g. extinction and re-acquisition during reversal learning) has been investigated with the PER assay as well (Eisenhardt and Menzel, 2007; Mota and Giurfa, 2010). Extinction describes the reduction in response to a previously learned stimulus when it is repeatedly presented without reward (Eisenhardt and Menzel, 2007). Reversal learning, on the other hand, consists in relearning the contingencies of stimuli (Mota and Giurfa, 2010). Extinction and reversal learning share common characteristics in that a previously formed association needs to be changed. They also both require processing in the mushroom bodies (MBs), a higher order brain center of bees (Devaud et al., 2007; Devaud et al., 2015).

Epigenetic mechanisms are crucial for transcriptional regulation (Rothbart and Strahl, 2014; Schubeler, 2015). They comprise mechanisms which can tightly and subtly regulate transcription, thus being good candidates for regulating complex behaviours. In bees, epigenetic mechanisms - such as histone acetylation and DNA methylation - have been related to memory formation (Biergans et al., 2016; Biergans et al., 2015; Biergans et al., 2012; Lockett et al., 2010; Merschbaecher et al., 2012). Following olfactory reward conditioning proteins catalyzing DNA methylation (Dnmts) and demethylation (Tet) are upregulated and DNA methylation levels change in memory-associated genes (Biergans et al., 2015). In the presence of a Dnmt inhibitor global DNA methylation levels decrease in the brain and memory-associated genes are upregulated 24 hours after training (Biergans et al., 2015). Furthermore, DNA methylation mediates associative plasticity in the neural network of the primary olfactory center and aids odour discrimination (Biergans et al., 2016). These studies support earlier behavioural data arguing for a role of DNA methylation in stimulus-specific LTM formation (Biergans et al., 2012) and extinction (Lockett et al., 2010).

Here we investigated in detail the behavioural phenotypes these studies describe. Specifically, we assessed whether the observed effects after Dnmt inhibition are learning-dependent, replicable and robust. Furthermore, we re-analysed all Dnmt inhibition experiments present to date to determine which functions of Dnmts during memory formation are best supported by the data.

## 2 Results

### 2.1 Dnmts do not affect odour or sugar perception in the absence of learning.

DNA methylation allows the animal to form a stimulus-specific LTM memory: when Dnmts are blocked, animals generalize more after learning. A possible explanation could be that Dnmt inhibition affects plasticity in stimulus perception rather than memory formation. Previous experiments already approached this hypothesis by treating bees with a Dnmt inhibitor 24 hours before training. In these experiments acquisition, memory retention and generalization did not change (Biergans et al., 2012; Lockett et al., 2010). Here, we confirm that Dnmts do not affect perception in a context without learning. We treated bees with the Dnmt inhibitor RG108 or the solvent DMF and tested their naïve odour preference (Fig. 1A) and sugar sensitivity (Fig. 1B) 22 hours later. Additionally, we used unpaired conditioning where bees received odour and sugar separated by 5 min (Fig. 1C). In this paradigm no memory is formed (Hellstern et al., 1998), but the cumulative stimuli experienced by the bee are the same as in appetitive learning studies. In all cases - odour preference, sugar sensitivity and unpaired conditioning - there was no difference between Dnmt inhibitor treated and control bees (odour preference: glm, factor treatment: hexanol: p=0.802; nonanol: p=0.409; hexanone: p=0.577; heptanone: p=0.156; sugar sensitivity: glm, factor treatment: p=0.314; unpaired conditioning: glm, factor treatment: CS: p=0.118; new: p=0.096). Thus, exposure to olfactory or gustatory stimuli in the absence of learning does not lead to DNA methylation changes that affect later odour responses. We conclude that the generalization effect observed after Dnmt inhibition is likely to be learning-dependent.

**Figure 1:**
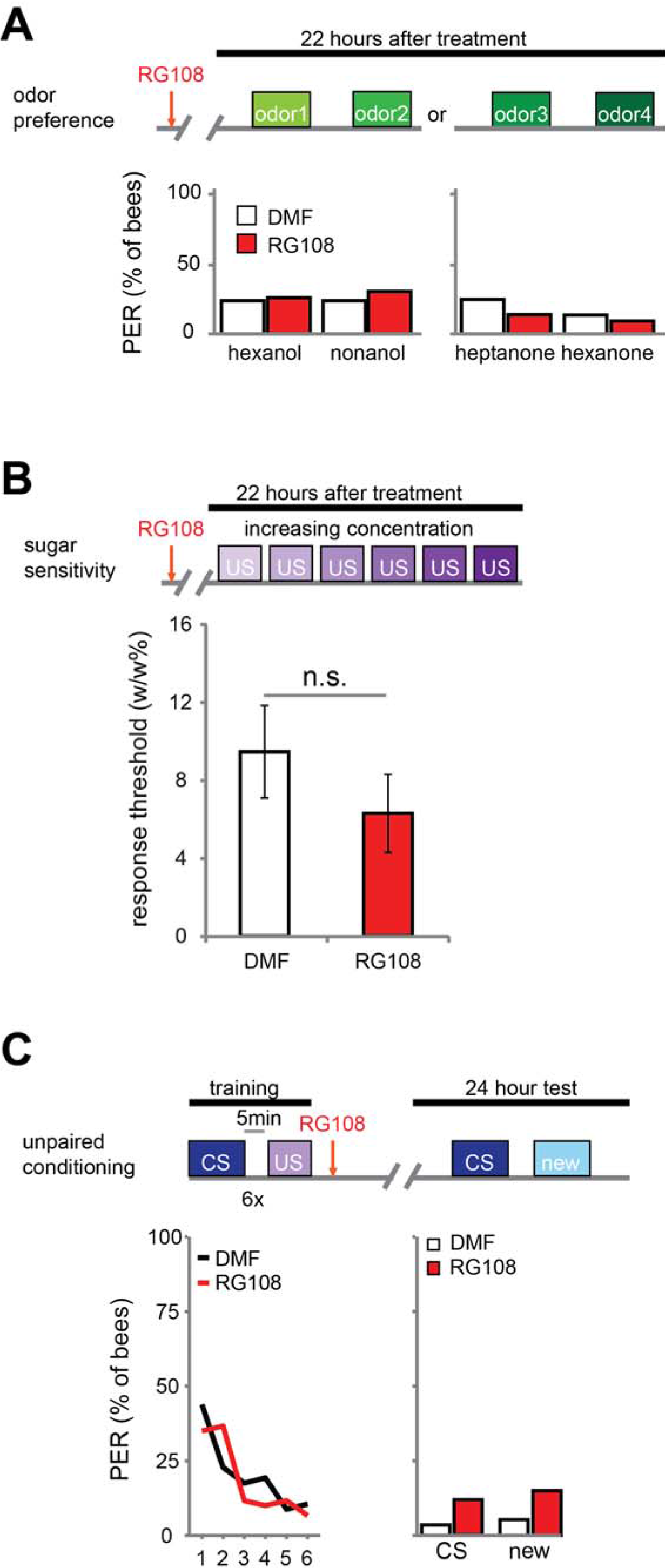
Dnmts do not affect odour or sugar perception in the absence of learning. (A) The percentage of bees naïvely responding to all odours used in this study is shown. Bees were treated with 1 μl of the Dnmt inhibitor RG108 or the solvent DMF 22 hours before the test, but no training took place. Two experiments: one with hexanol and nonanol (n(RG108)=65, n(DMF)=58), one with heptanone and hexanone (n(RG108)=43, n(DMF)=44). naïve odour responses were not different after RG108 treatment; (glm, factor treatment: hexanol: p=0.802; nonanol: p=0.409; hexanone: p=0.577; heptanone: p=0.156). (B) Bees were tested for their sugar responsiveness 22 hours after RG108 treatment. Increasing concentrations of sugar water (0.1 - 30 %w/w) were presented to their antennae. The response threshold is shown (mean +/− SEM). The response threshold was not different between RG108 and solvent treated bees (n(DMF)=28, n(RG108)=27; glm, p=0.314). (C) Although naïve odour responses were not affected by Dnmt inhibition the pre-exposure to the stimuli during training could be sufficient to change the response in the test even in the absence of learning. To control for a possible effect of pre-exposure we trained bees with an unpaired paradigm (5 min between the CS and US), treated them with RG108 or the solvent 2 hours after training, and tested their response to the pre-exposed and a new odour 22 hours later. The response did not differ between treatments (n(RG108)=60, n(DMF)=57; glm, factor treatment: CS: p=0.118; new: p=0.096).

### 2.2 Methylation adjusts the strength of generalization depending on the training conditions.

Next, we investigated which training parameters influence how Dnmts affect stimulus-specific memory. We utilised two variations of PER conditioning which initiate distinct molecular pathways: single-trial learning, and multiple-trial learning. First, we tested one trial training (i.e. only one odour-sugar pairing, Fig. 2A). Control bees had a weak stimulus-specific memory after 24 hours (Fig. 2B). After Dnmt inhibition, however, bees formed a stimulus specific memory, successfully discriminating between the CS+ and a new odour (McNemar test, p=0.011, effect size=0.32). The number of bees responding correctly only to the CS+ increased after RG108 treatment (Fig. 2C, chi^2^-test, p=0.014, effect size=0.37). Thus, after one-trial-training, Dnmt activity reduced odour selectivity in the memory trace. Next, we tested multiple-trial (massed) training. We trained bees with 6 odour-sugar pairings, separated by 1 minute each (Fig. 2D). When Dnmts were inhibited, stimulus-specific memory formation was impaired and discriminatory power was significantly lower compared to control bees (Fig. 2E, glm, p=0.008, effect size=0.56). Both the number of bees responding 'correctly' only to the CS+ was reduced (Fig. 2F, chi^2^-test, p=0.008, effect size=0.56), and the number of bees responding 'wrongly' to both test odours was increased after Dnmt inhibition (Fig. 2F, chi^2^-test, p=0.026, effect size=0.46). These data supplement previously published data with spaced multiple trial training (10 min intertrial interval), which also showed increased generalization when DNMTs were blocked (Biergans et al., 2015; Biergans et al., 2012). Thus, while DNA methylation increases generalization after one trial learning, DNA methylation decreases generalization (increases odour recognition) in multiple-trial learning, leading to a more selective odour response (Biergans et al., 2016). This is an intriguing bi-directional effect of DNA methylation.

**Figure 2:**
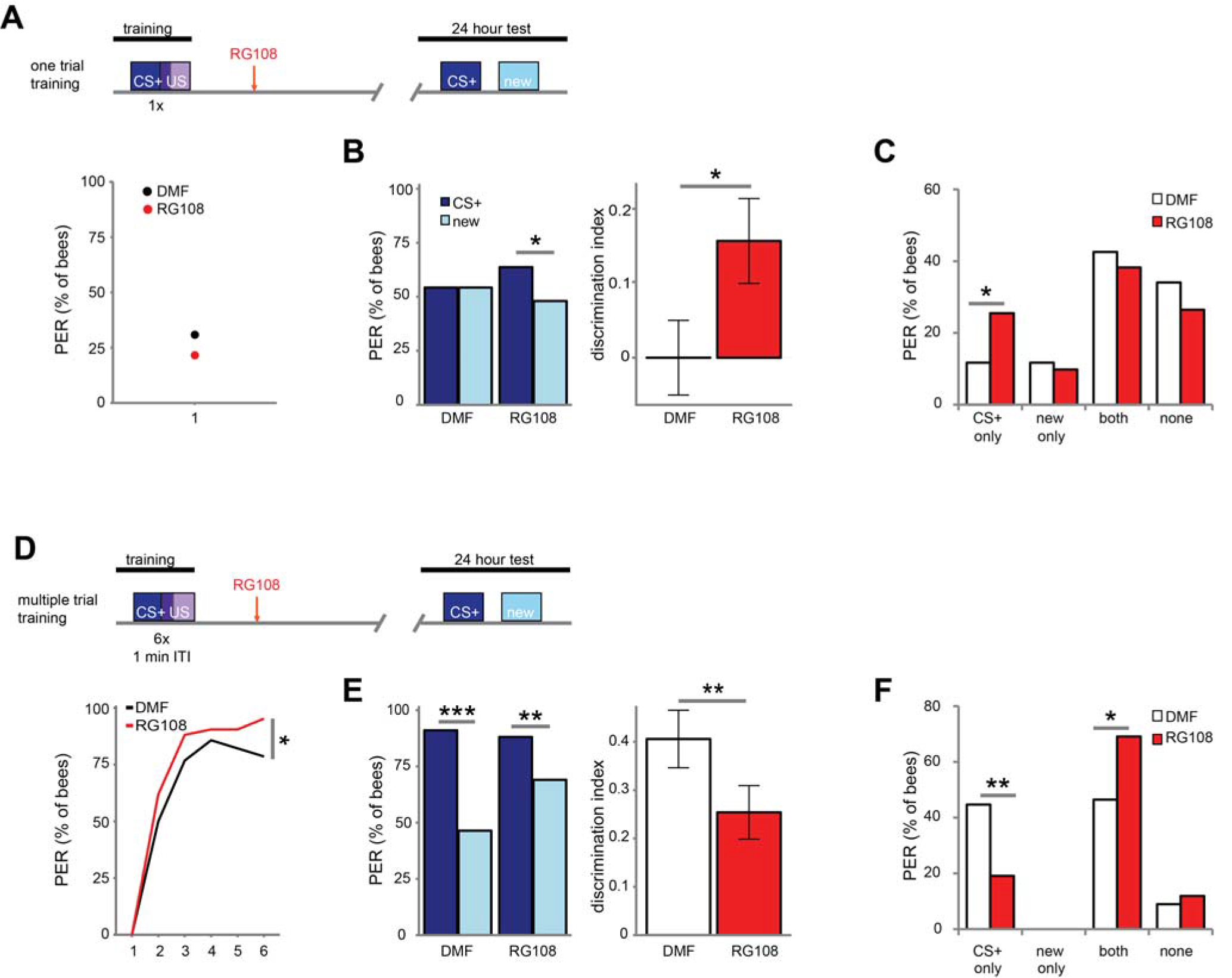
Dnmts influence stimulus-specific memory bidirectionally. ((A) Bees were trained with one CS-US pairing (D) or six CS-US pairings with an inter-trial interval (ITI) of 1 minute. 2 hours after the training bees were treated with the Dnmt inhibitor RG108 or the solvent DMF and tested for memory retention (CS+ response) and generalisation (new response) after 24 hours. (B) Solvent treated bees did not show stimulus-specific memory in the 24 hours test following one CS-US pairing, but bees were able to discriminate between the CS+ and new odour after Dnmt inhibition (n(DMF)=94, n(RG108)=102, PER CS-new: McNemar test, p=0.011, effect size=0.32; discrimination index: glm, p=0.042, effect size=0.29). (C) Bees were sorted into responding groups: bees responded more often only to the CS+ after Dnmt inhibition (chi^2^-test, p=0.014, effect size=0.37). (E) Bees’ stimulus-specific memory was impaired in RG108 treated bees after multiple trial training (n(DMF)=56, n(RG108)=42; glm, p=0.008, effect size=0.56) (F) Bees responded less to the CS+ only (chi^2^-test, p=0.008, effect size=0.56) and more often to both odours (Chi^2^-test, p=0.026, effect size=0.46).

### 2.3 Dnmts regulate both extinction and re-acquisition.

Dnmts are also involved in extinction learning and memory (Lockett et al., 2010); i.e. the reduced response to a previously learned odour ('*extinction*') when the odour is repeatedly given without reward. We investigated whether Dnmts are also involved in relearning a previously forgotten odour ('*re-acquisition*')? We used a reversal learning paradigm. We trained bees three times, where each training was separated by 24 hours (Fig. 3A). Training was differential with one rewarded (CS+) and one unrewarded odour (CS−). The contingencies of odours were reversed every day, meaning that the odour which was rewarded on day one and three was unrewarded on day two, and vice versa.

**Figure 3:**
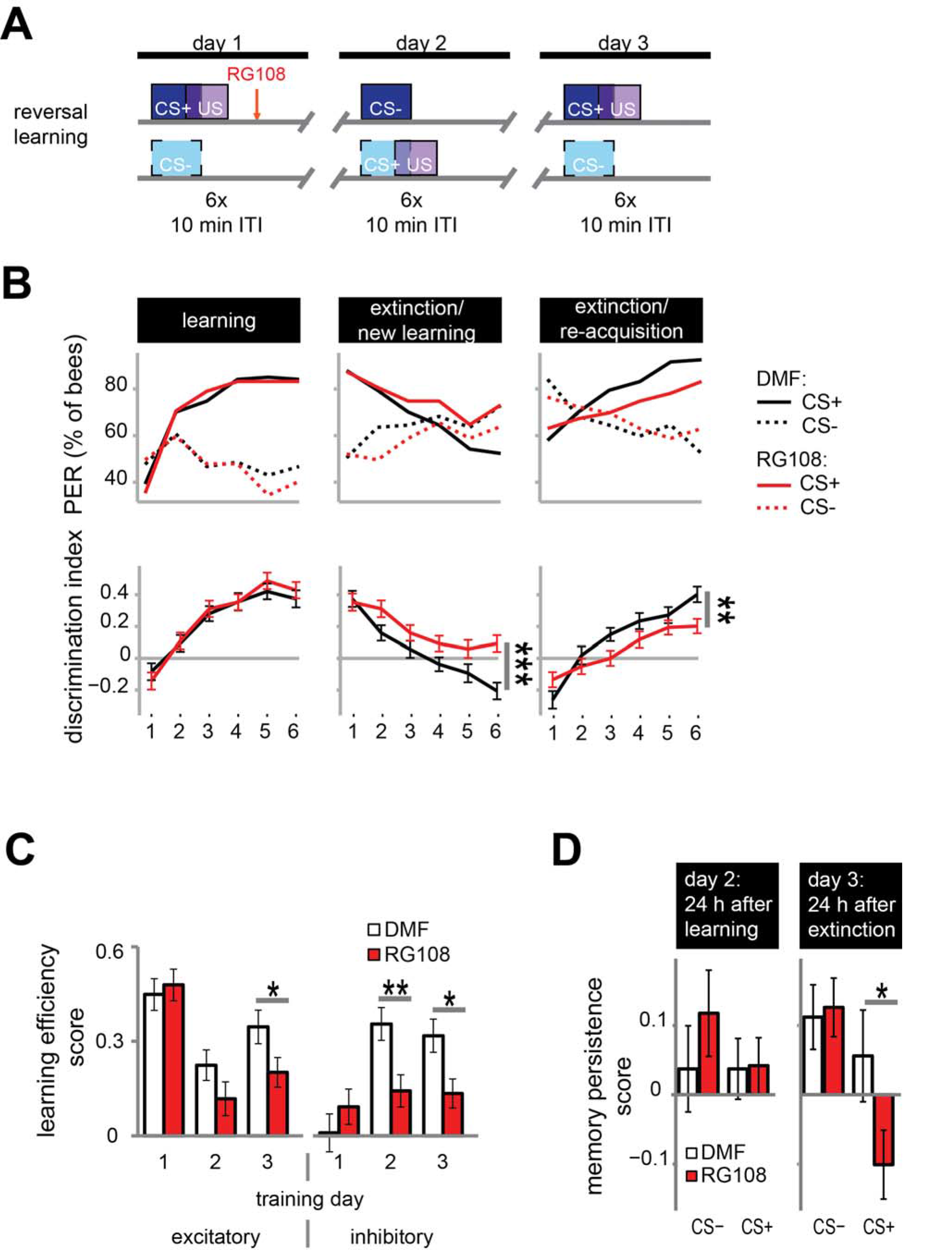
Dnmts promote the relearning of previously learned stimuli. (A) Bees were trained with a reversal training paradigm. (B) Solvent treated control bees responded correctly at the end of each training. RG108 treated bees, however, did not reverse the odour contingencies on day two (extinction/new learning) and performed significantly worse than control bees (n(DMF)=107, n(RG108)=119, glm, p<0.001, effect size=0.54). On day three (extinction/re-acquisition) they learned the reversal but did so significantly slower than control bees (glm, p=0.005, effect size=0.40). (C) Learning efficiency scores (bees’ last minus its first training trial response; 0 = same response, 1 = successful learning, −1 = opposite response) were calculated for the excitatory and inhibitory components in each training. The excitatory component was impaired by RG108 treatment only on training day 3 (glm, p=0.050, effect size=0.27). The inhibitory component was impaired on training day 2 and 3 (glm, day 2: p=0.004, effect size=0.39, day 3: p=0.013, effect size=0.35). (D) Memory persistence scores (difference in a bees’ response between the last training trial and the first one 24 hours later; 0 = same response, 1 = increase, −1 = decrease) were calculated for each period between trainings. RG108 treatment did not change the response to the CS+ or the CS− over the 24 hours initial learning. During the second 24 hours, however, RG108 treated bees changed their response to the unrewarded odour, but control bees did not (glm, p=0.032, effect size=0.26).

Control bees showed strong learning on day one, extinction and new learning within four trials on day two, and extinction and re-acquisition within three trials on the third day (black lines in Fig. 3B). When treated with a Dnmt inhibitor (red lines in Fig. 3B), however, bees were not able to learn the reversed contingencies of the odours on day two, performing significantly worse than control bees (glm, p<0.001, effect size=0.54). Dnmt-inhibited bees were also significantly slower in learning during the extinction/re-acquisition phase on day three compared to control bees (glm, p=0.005, effect size=0.40).

Reversal learning consists of two components - an excitatory (i.e. increasing the response to the previously unrewarded odour) and an inhibitory component (i.e. decreasing the response to the previously rewarded odour) (Mota and Giurfa, 2010). Thus, we analysed these components separately in order to investigate whether Dnmts are involved in the regulation of either or both. We calculated the learning efficiency score for each training day and stimulus by subtracting the bees’ response in the first training trial from its response in the last (Fig. 3C: 0 = no change in response, 1 = show learned response, −1 = show opposite effect) as described elsewhere (Mota and Giurfa, 2010). Dnmt inhibition caused a reduction of the inhibitory component on training day 2 and 3 and of the excitatory component on training day 3 (Fig. 3C; glm, excitatory: day 2: p=0.050, effect size=0.27; inhibitory: day 2: p=0.004, effect size=0.39, day 3: p=0.013, effect size=0.35). Thus, both extinction (i.e. inhibitory component) and re-acquisition (i.e. excitatory component) relied on DNA methylation.

Next, we investigated whether the response after memory consolidation was also affected by the treatment induced impairments observed during training. We calculated a memory persistence score by subtracting the bees’ response in the last training trial from its response in the first training trial 24 hours later (Fig. 3D: 0 = same response 24 hours after training, 1 = increased response, −1 decreased response). The bees’ responses at the end of the learning phase on day 1 were largely maintained at the beginning of the day 2 (Fig. 3D '24 h after learning'). 24 hours following the extinction/new learning phase, however, bees’ memory retention was improved for the initial CS+. RG108 treated bees also maintained the response to the initially learned odour at the beginning of day 2, but showed reduced responses to that odour at the beginning of day 3 (Fig. 3D '24 h after extinction', glm, p=0.032, effect size=0.26). This suggests that the impairment of the inhibitory learning component during the 2^nd^ training day is compensated during memory consolidation between days 2 and 3.

The necessity of active Dnmts during reversal learning could indicate that Dnmts are either important for the re-learning of previously learned stimuli or for the ability to learn in general. So far there is more evidence for the first hypothesis, as Dnmts are not necessary during acquisition in naïve bees (Biergans et al., 2012; Lockett et al., 2010). In the paradigm used in Fig. 3, however, bees learned in the context of previous training, creating a situation where the effect of learning ability or re-learning ability cannot be separated. Therefore, we modified the protocol as follows: We trained bees as described before with a differential training paradigm including a rewarded (CS+) and an unrewarded odour (CS−). On day two however - instead of re-training with the previously used odours - we trained bees with two new odours (Fig. 4A). We found that both the solvent treated control bees and RG108 treated bees were able to learn to discriminate the new odours during the 2^nd^ training day (Fig. 4B). None of the learning components was affected by Dnmt inhibition (Fig. 4C). This confirms that Dnmt activity was not important for acquisition in general, but it was important specifically for the relearning of previously learned stimuli.

**Figure 4:**
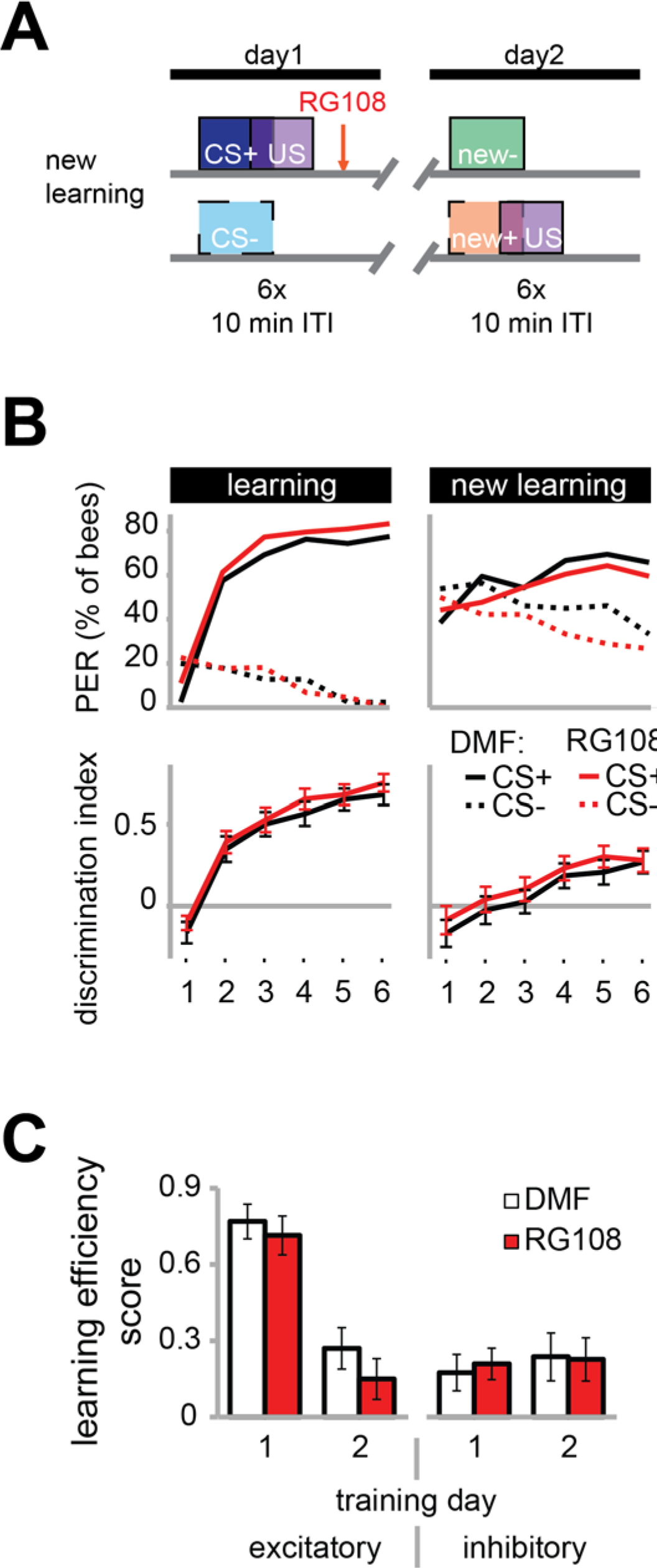
Dnmts do not regulate the learning of new odours. ((A) To confirm that Dnmts are necessary for efficient relearning of previously learned stimuli, but not for acquisition in general, we trained bees with a differential conditioning paradigm as in Fig. 3, but trained them with two new odours on day two. (B) Bees learned to discriminate the two new odours with and without active Dnmts. (C) Neither excitatory nor inhibitory components of learning were impaired by Dnmt inhibition on the second training day. n(DMF)=39, n(RG108)=44

### 2.4 A role of Dnmts in 'correct' LTM formation, reversal learning and extinction MTM is supported by the experimental data to date.

In this study, we presented experiments showing that Dnmts regulate stimulus-specific LTM and re-learning, but do not affect stimulus perception or acquisition of new stimuli. In order to compare these results to and imbed them with the body of data available in the literature so far, we performed a meta-analysis. We aggregated available published data (Biergans et al., 2015; Biergans et al., 2012; Lockett et al., 2010), unpublished data (summarised in Supplemental Tab. 1) and all experiments shown in this study. We formed three categories: (1) experiments testing LTM (Fig. 5A,B), (2) experiments testing re-learning (Fig. 5C,D) and (3) control experiments (Fig. 5E,F). All experiments used odour reward conditioning and PER as a behavioural read-out. They differed, however, in the training paradigm used (i.e. absolute or differential) and the stimuli tested (CS+, CS−, new odour, sugar). We calculated the % of bees responding ‘correctly’ in the inhibitor treated group (reduced Dnmt activity) of each experiment and plotted this value against the % of bees responding correctly in the solvent treated group (normal Dnmt activity, Fig. 5A,C,E). The scores differed across the training/test paradigm used (summarised in: Tab. 1). The individual experiments further differed in the inhibitor used (i.e. RG108 (**X**) or Zebularine (**O**)) and the treatment timepoint (i.e. before (yellow: **O**), before+after (green: **O**) or after (blue: **O,X**) training). In Fig. 5A, a point on the diagonal indicates an experiment where Dnmt activity does not affect‘correct’ LTM formation; points below the diagonal (lower-right) indicate experiments where Dnmt activity positively contributed to ‘correct’ memory performance.

**Figure 5:**
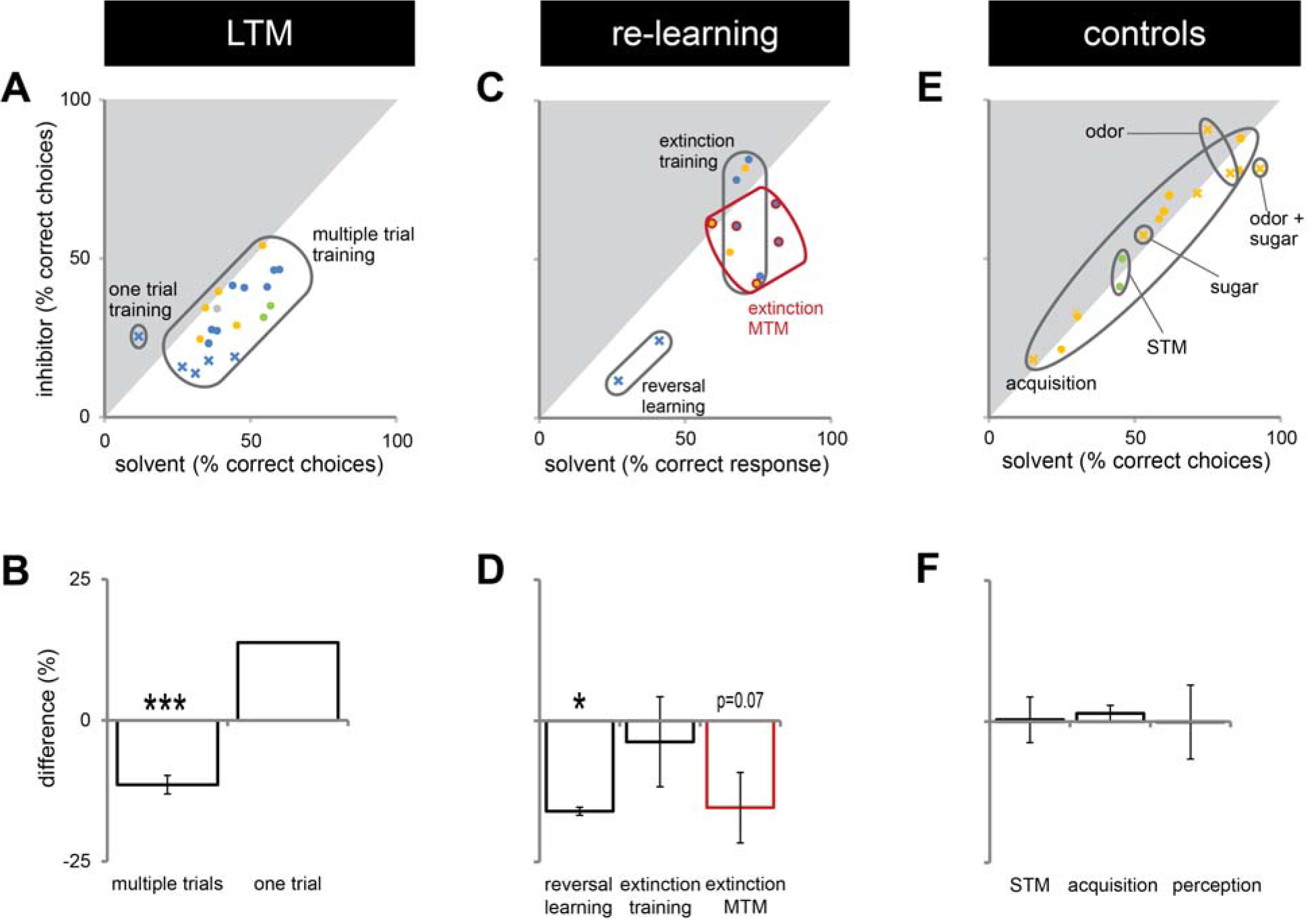
Correct LTM formation and relearning are both facilitated by Dnmts. All PER experiments shown in this study, published previously (Biergans et al., 2015; Biergans et al., 2012; Lockett et al., 2010) or performed additionally (summarised in Supplemental Tab. 1), were re-analysed in order to gain an overview of which roles of Dnmts are best supported by the data. The % of bees responding 'correctly' was calculated for the treatment and control groups. What a 'correct' response was differed between experiments (Tab. 1). (A,C,E) The % of correct responses for the solvent and inhibitor treated groups are plotted against each other. Each mark represents one experiment (inhibitor: X=RG108, O=zebularine; treatment time-point: yellow=before; green=before+after; blue=after training). (A) Solvent treated bees responded 'correctly' more often than inhibitor treated bees in most experiments testing LTM retention after multiple trial olfactory reward conditioning. (B) Pooling the difference between the ‘correct’ responses in inhibitor and solvent treated bees of all experiments shows a significant effect of Dnmt activity (n=21, one sample t-test, p<0.001, effect size=1.516). (C,D) Reversal learning (n=2, one sample t-test, p=0.028, effect size=16.052) and extinction MTM (n=5, one sample t-test, p=0.070) were most consistently impaired by Dnmt inhibition, whereas the results for extinction learning (n=5, one sample t-test, p=0.665) were inconsistent. (E,F) Control experiments so far tested the effect of Dnmt inhibition on STM, acquisition and perception. There was no consistent effect on either (n(STM)=2, n(Acquisition)=10, n(Perception)=4; one sample t-test, STM: p=0.951, Acquisition: p=0.343, Perception: p=0.990).

Most LTM experiments showed a reduction of 'correct' responses after Dnmt inhibition (points below the diagonal in Fig. 5A,B, one sample t-test, p<0.001, effect size=1.516). Thus, the data suggest a role of Dnmts in facilitating 'correct' LTM formation with an average reduction of 11% in ‘correct’ responses when Dnmts are pharmacologically inhibited. Data also show a role of Dnmts in extinction mid-term memory (MTM, Fig. 5C,D, red circles, one sample t-test, p=0.070) and reversal learning (Fig. 5C,D, one sample t-test, p=0.028, effect size=16.052). For extinction learning, however, data are inconsistent, with some experiments showing a reduction in extinction and some an increase after Dnmt inhibition (Fig. 5C,D, one sample t-test, p=0.665). The difference may be due to the treatment time-point. In order to control for a potential effect of Dnmts on acquisition or stimulus perception, all control experiments performed were also pooled (Fig. 5E,F). There was no consistent effect on short-term memory, acquisition or perception.

## 3 Discussion

Long-term memory needs neural networks that are modified in a stable way over several days up to a life time. Epigenetic modifications of the genome in neurons have been shown to contribute to these long-term memory traces (Zovkic et al., 2013). However, every memory consists of several phases (over time), content (e.g. stimulus specificity, associative strength), and involves distinct neural networks across the brain. Therefore, it is important to dissect the exact role of a particular molecular mechanism in order to build a complete picture of how memory functions in a living animal. Here, we show that DNA methylation fulfils very specific roles in a honeybee olfactory reward memory paradigm. Specifically, we show that Dnmts regulated stimulus-specific long-term memory robustly under differing training parameters, whereas the directionality of the regulation depended on the trial number during training (Fig. 2). Additionally, we show that Dnmts regulated extinction and thus the inhibitory component of re-learning, and also the excitatory component in a reversal learning paradigm (Fig. 3). Furthermore, we re-evaluated the evidence available to date focusing on the role of Dnmts in olfactory reward conditioning and found that Dnmts consistently play a role in 'correct' long-term memory formation (e.g. stimulus-specific memory formation), reversal learning and extinction MTM, whereas its role in extinction learning remains unresolved at this point (Fig. 5).

Dnmt inhibition did not affect naïve odour and sugar responses or odour responses after unpaired training (i.e. without memory formation). These results confirm earlier studies showing that acquisition is not affected by Dnmt inhibition 24 hours before (Biergans et al., 2012; Lockett et al., 2010), which argues against an effect of Dnmts on naïve stimulus perception. This suggests that Dnmt activity (and/or expression) is induced by the coincident occurrence of CS and US during learning. Indeed, it has recently been shown that Dnmts and also the demethylation protein Tet are upregulated after olfactory reward conditioning (Biergans et al., 2015).

DNA methylation via Dnmts regulates stimulus-specific LTM after olfactory reward conditioning (Biergans et al., 2015; Biergans et al., 2012). The directionality of this regulation, however, depended on the number of training trials, and was independent of the inter-trial interval (Fig. 2). After one odour-sugar pairing control bees formed a weak stimulus-specific memory, which confirms earlier studies (Lefer et al., 2012; Perisse et al., 2009). Following Dnmt inhibition, however, bees were able to discriminate between the trained and a new odour. Thus, Dnmt-dependent mechanisms seem to increase generalization after one trial training and decrease generalization after multiple trial training. Without the activity of Dnmts, generalization is comparable in the two situations. Thus, we can speculate that the adaptive role of Dnmts in regulating memory is the following: a single odour-sugar pairing is not sufficient to predict that a particular odour is rewarded, and Dnmt activity therefore weakens the odour identity related information in the memory trace. Repeated pairings, on the other side, indicate reliable odour-information, and methylation increases odour-specific memory information. This differential effect may be based on differing molecular pathways. Multiple trial training induces long-lasting PKA and PKC activity and is counter-acted by protein degradation, whereas one trial training is not (Felsenberg et al., 2012; Grünbaum and Müller, 1998; Hildebrandt and Muller, 1995; Muller, 2000). At this point there is not enough information about how Dnmts may regulate stimulus-specific memory bidirectionally. We, however, discuss two tentative possibilities: (1) Dnmts have been shown to have demethylating activity, in addition to their predominant methylating activity (Chen et al., 2013). This reversal in function is related to Ca^2+^ levels. Differences in stimulus-specific memory formation between one and multiple trial training may be related to differing Ca^2+^ levels after training (Perisse et al., 2009). Thus, different Ca^2+^ levels present in neurons after training may regulate whether Dnmts are active as methylase or as demethylase. It has to be noted though that Dnmt demethylase activity has only been described under specific conditions *in vitro* yet. (2) Another possibility is that the different molecular pathways triggered after one and multiple trial training (Felsenberg et al., 2012; Hildebrandt and Muller, 1995; Muller, 2000; Perisse et al., 2009) cause Dnmts to target different genes. Similarly, histone modifications also follow different dynamics after one and multiple trial training (Merschbaecher et al., 2012). Thus, Dnmts and related up-and downstream processes might fine-tune memory formation depending on the environmental information available and thus allow for maximally beneficial adjustments in an animals’ behaviour.

Dnmts also regulate extinction in bees (Lockett et al., 2010). Extinction is a form of relearning during which bees need to re-evaluate a previously rewarded stimulus as being not rewarded any more. Compared to this, during reversal learning, bees have to re-evaluate two stimuli simultaneously, with one being rewarded and the other one not. Evidence from both extinction and reversal learning studies favors the idea that 'positive' and 'negative' memories of a stimulus are present in parallel (Mota and Giurfa, 2010; Stollhoff et al., 2005). Our data suggests that both the inhibitory component of reversal learning (i.e. extinction) and the excitatory component (i.e. re-acquisition) involve Dnmts. Dnmts may regulate these processes in two ways: (1) Dnmts could affect the balance between opposing memory traces for a stimulus. This could cause a behavioural dominance of the most recent association learned over older memories during training. (2) Dnmts could be involved in de-constructing the older memory trace. Further experiments investigating extinction and reversal learning and the underlying molecular mechanisms are needed to gain insight into what the specific function of Dnmts is here. Interestingly, even though relearning was impaired during the training, memory recall 24 hours after training was not, whereas subsequent relearning was again impaired. Thus, it seems as if Dnmts regulate pathways needed during relearning, but that the memory is consolidated correctly without the need for DNA methylation. Notably, only the relearning of previously learned stimuli was impaired when Dnmts are inhibited, but not bees’ general ability to learn. This suggests that Dnmts set methylation marks only in those neurons active during training (e.g. neurons responding to a particular odour) and thus potentially create a memory trace on the level of the chromatin mirroring the activity of that neuron over time.

With 'correct' LTM formation and relearning, we now know that Dnmts are involved in two distinct groups of behavioural readouts after olfactory reward conditioning in bees (Fig. 4). Further studies will need to investigate how exactly Dnmts regulate these behaviours and whether the same Dnmt targeted genes affect both, or whether distinct sets of genes are required for each. Furthermore, it will be crucial to investigate the role Dnmts play in learning paradigms utilizing different CS (e.g. visual stimuli) and US (e.g. punishment). This will reveal which neuronal networks and brain centers are involved and whether Dnmts regulate the same processes independent of the sensory modalities utilized during training.

## 4 Materials and Methods

### 4.1 Odour reward conditioning and memory retention test

Experiments were performed either at the University of Queensland (Brisbane, Australia) or the University of Konstanz (Konstanz, Germany). Honey bees (*Apis mellifera*) were caught outside the hive and put on ice until they were immobilized. Bees were harnessed in plastic tubes so that they could only move their head, but with their thorax still accessible. They were fed until satiation and kept overnight in a humid plastic box, or an incubator depending on where the experiment was performed. The next day bees were trained using an appetitive olfactory training paradigm. The exact training parameters were different for each experiment (summarised in Tab. 2). In all experiments the odour was presented for 4 s and sugar reward (1 mM sugar water) for 3 s. Odours (all Sigma-Aldrich) were dissolved in hexane for the experiments performed in Australia and in mineral oil (in all cases diluted 10^2^) for those performed in Konstanz.

**Table 2.** Overview over training parameters. For each experiment presented here the number of trials (NoT), inter-trial interval (ITI), inter stimulus interval (ISI) and location, where the experiment was performed, is shown.

| Training/Test | CS+ | CS- | New odour | Sugar |
| --- | --- | --- | --- | --- |
| Differential training/test | PER | No PER | X | X |
| Absolute training/test | PER | X | No PER | X |
| Unpaired training/test | No PER | X | No PER | X |
| Extinction learning/MTM | No PER | X | X | X |
| Naïve odour test | X | X | No PER | X |
| Naïve sugar test | X | X | X | PER |

During each experiment bees experienced a constant air-flow in order to avoid mechanical stimulation at odour onset. 24 hours after training bees were tested for memory retention by presenting them with the CS+ (trained odour) and a new odour, which was not present during the training, in randomised order. 1-hexanol and 1-nonanol were alternated as CS+, CS− or new odour respectively. Additionally, for the control experiments shown in Fig. 4 1-heptanone and 1-hexanone were used on the second day for training.

### 4.2 Control experiments

During the control experiments investigating whether Dnmt inhibition affects stimulus perception (Fig. 1), bees did not receive olfactory appetitive training. Instead, bees were tested for ‘naïve’ odour or sugar responses 22 hours after treatment - equivalent to the time trained bees were tested after treatment. For the odour preference test, bees were tested for their spontaneous proboscis extension response to all 4 odours used here in two separate experiments. 1-hexanol and 1-nonanol were always tested together and their order was alternated across bees (the same for 1-hexanone and 1-heptanone). For their sugar response bees were tested with increasing concentrations of sugar water (0.1, 0.3, 1, 3, 10, 30 %w/w). Bees’ antennae were touched with a tooth pick soaked in sugar water and it was recorded whether or not bees extended their proboscis in response. The lowest concentration a bee responded to (response threshold) was compared between treatments. Before and after the test, as well as after each individual sugar concentration, bees were tested for their response to water. Bees responding to water more than twice or not responding to the highest sugar concentration were discarded from the experiment.

### 4.3 Treatment

2 hours after training and 22 hours before the control experiments bees were treated with 1 Ml of the Dnmt inhibitor (RG108, 2 mM in DMF, Sigma-Aldrich) or the solvent DMF on the thorax as described elsewhere (Biergans et al., 2015; Biergans et al., 2012; Lockett et al., 2010).

### 4.4 Data analysis

For all experiments the % of bees responding to the odours in the test and training was calculated. Furthermore, a discrimination index was calculated. The response of each individual bee to the new odour or CS-was subtracted from its response to the CS+. All data were analysed using generalised linear models, if treatment groups were compared. To compare the response to the CS+ and new odour within one treatment group, a McNemar test was used. The McNemar test is suitable to compare binary, paired data.

For the meta-analysis we gathered all honeybee data investigating the role of Dnmts in memory formation, including published data (Biergans et al., 2015; Biergans et al., 2012; Lockett et al., 2010), data presented in this study and unpublished data. An overview over all data sets used for the meta-analysis is shown in Supplemental Tab. 1. We calculated the number of 'correct' responses within each experiment and experimental group (Tab. 1).

**Table 1:**
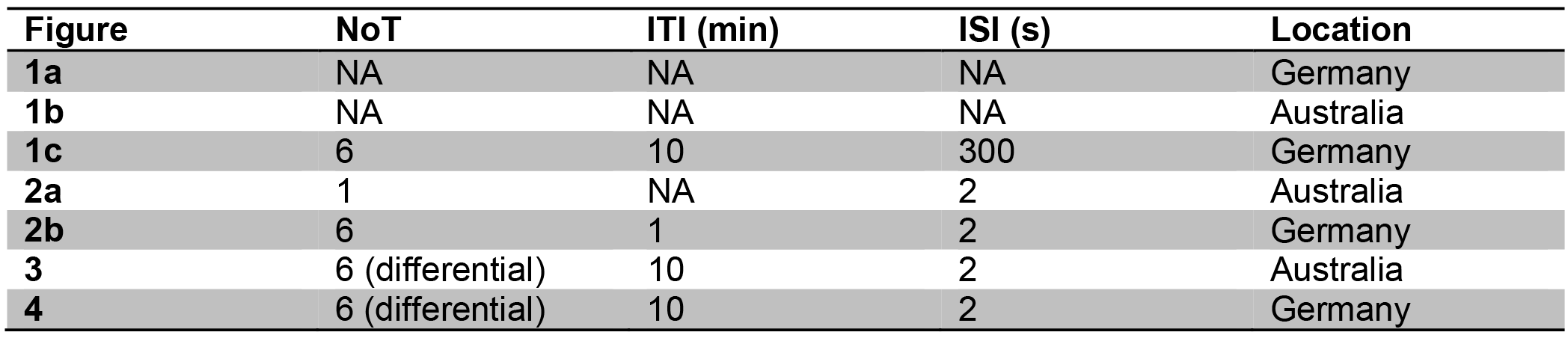
Evaluation of correct behavioural responses in the analysed data sets. Depending on the training paradigm and test used a ‘correct’ response differed. The trained and tested stimuli are shown, and what comprised a ‘correct’ response in the specified training/test. ‘X’ indicates that this stimulus was not present in the specified training/test.

Using this method we were able to compare data obtained by different training paradigms and assess the overall evidence for the effect of Dnmt inhibition. To quantify the level of agreement between different studies we calculated the difference in the correct responses of inhibitor and solvent groups for each experiment and pooled them. A two-sided one-sample t-test was used to test whether the effect shown in those studies is reliably different from 0. The effect size (Cohen's D, (Navarro, 2015)) was calculated for all effects reaching the 0.05 significance level. As a guideline effect sizes below 0.2 are described as negligible, between 0.2 - 0.5 as small, between 0.5 - 0.8 as medium and above 0.8 as large (Cohen, 1992). The effect size can be used as an estimate of the real difference between the tested groups. All analyses were performed using custom written R-scripts using R-version 3.2.1 (R Core Team, 2015).

## Acknowledgments

We thank Ryszard Maleszka and colleagues for sharing their data with us. We thank Ulrike Schlegel and Sabrine Kurzeja for helping with data collection. We also thank Morgane Nouvian for helpful feedback for the manuscript. The study was funded by grants from the Australian Research Council and National Health & Medical Research Council of Australia (ARC DP120104117, ARC DP120102301, NHMRC APP1008125 to CC and JR), and by the Deutsche Forschungsgemeinschaft DFG (SPP 1392 to CGG). CC was also supported by an Australian Research Council Future Fellowship (FT110100292). SDB was funded by a University of Queensland Postgraduate Research Scholarship and travel award.

## Author contributions

SDB, CC, JR and CGG conceived and designed the study. SDB performed the experiments. SDB analysed the experiments. SDB wrote the paper. CC and CGG edited the paper.

